# The Proteolytic Landscape of an Arabidopsis Separase-Deficient Mutant Reveals Novel Substrates Associated With Plant Development

**DOI:** 10.1101/140962

**Authors:** Chen Liu, Simon Stael, Kris Gevaert, Frank Van Breusegem, Peter V Bozhkov, Panagiotis N Moschou

## Abstract

Digestive proteolysis executed by the proteasome plays an important role in plant development. Yet, the role of limited proteolysis in this process is still obscured due to the absence of studies. Previously, we showed that limited proteolysis by the caspase-related protease separase (EXTRA SPINDLE POLES [ESP]) modulates development in plants through the cleavage of unknown substrates. Here we used a modified version of the positional proteomics method COmbined FRActional DIagonal Chromatography (COFRADIC) to survey the proteolytic landscape of wild-type and separase mutant *RADIALLY SWOLLEN 4* (*rsw4*) root tip cells, as an attempt to identify targets of separase. We have discovered that proteins involved in the establishment of pH homeostasis and sensing, and lipid signalling in wild-type cells, suggesting novel potential roles for separase. We also observed significant accumulation of the protease PRX34 in *rsw4* which negatively impacts growth. Furthermore, we observed an increased acetylation of N-termini of *rsw4* proteins which usually comprise degrons identified by the ubiquitin-proteasome system, suggesting that separase intersects with additional proteolytic networks. Our results hint to potential pathways by which separase could regulate development suggesting also novel proteolytic functions.

## Introduction

Separase (family C50, clan CD) is an evolutionary conserved cysteine protease participating in the metaphase-to-anaphase transition and plant development (Liu & Makaroff, 2006; Moschou & Bozhkov, 2012). In *Arabidopsis thaliana*, separase localizes to microtubules (Moschou *et al*., 2016a), the plasma membrane and the endosomal compartment trans-Golgi Network (TGN)(Moschou *et al*., 2013), which acts as a hub for protein vesicle trafficking.

Since loss-of-function Arabidopsis separase mutants are embryo lethal (Liu & Makaroff, 2006), the roles of separase beyond cell division are studied by the conditional temperature-sensitive loss-of-function allele *RADIALLY SWOLLEN 4* (*rsw4*). The *rsw4* mutation renders separase unstable at 28^o^C (Wu *et al*., 2010; Yang *et al*., 2011; Moschou *et al*., 2016a). *Rsw4* shows root phenotypes reminiscent of compromised auxin distribution, with perturbed gravitropism, cell expansion and overall development (Moschou *et al*., 2013). Proteolytic activity of separase is indispensable in development since the proteolytic inactive separase mutant could not rescue *rsw4* defects. Yet, the proteolytic targets of separase remain obscure.

Implementation of positional proteomics is the most direct method for unbiased identification of protease substrates (Plasman *et al*., 2013). When sampling complex proteomes, protease substrates can be identified by exploiting the chemical reactivity of the alpha-amino groups. Such newly introduced α-amino groups are typically referred to as neo-N-termini (neo-Nt). The N-terminal COmbined FRActional DIagonal Chromatography technology (COFRADIC; (Gevaert *et al*., 2003) depletes non-Nt-peptides, thereby enriches for Nt-peptides and further allows the specific identification of neo-Nt through a combination of stable isotope labelling of neo-Nt, tandem mass spectrometry and bioinformatics.

Here we used COFRADIC to identify potential separase substrates that may allow delineating new signalling pathways relevant to organ growth and development. By contrasting the proteolytic landscape of wild-type (WT) and *rsw4* we infer potential proteolytic targets relevant to vesicular trafficking, lipid signaling and root growth control. We succinctly discuss the most relevant identified proteins and suggest potential molecular mechanisms for further exploration.

## Results and Discussion

### Inducible depletion of separase phenocopies rsw4 PIN2 polarity defects

A potent readout for vesicular trafficking defects and early events imposed by the absence of *rsw4* is the mis-localization of the auxin efflux carrier PINFORMED2 (PIN2) in root cortex cells, responsible for auxin flow maintenance (Moschou *et al*., 2013). Depletion of separase in *rsw4* causes PIN2 localization switch from the proper rootward to the shootward side of the plasma membrane in *rsw4* root cortex cells at the restrictive temperature. This PIN2 localization switch correlates with the reduced activity of a GNOM-related vesicular trafficking pathway in *rsw4*. A functional GNOM-related pathway requires proteolytic cleavage of an as yet unknown substrate(s) by separase, since the protease-dead allelic variant of separase could not complement PIN2 localization defects of *rsw4* (Liu & Moschou, 2017).

We have established that treatment of *rsw4* seedlings for up to 24 h at the restrictive temperature (28°C) disrupts separase activity and causes vesicular trafficking defects, but does not compromise the rate and pattern of cell division and the identity of root cells (Moschou *et al*., 2013). We argue that this experimental setup enables to study roles of separase beyond its mitotic role (Moschou *et al*., 2016b). Note that a temperature of 28°C falls within the physiological temperature range for Arabidopsis growth (Faden *et al*., 2016). Yet, to ensure that the *rsw4*-mediated effect in PIN2 polarity is not due to the temperature shift, we surveyed PIN2 polarity in dexamethasone inducible RNAi lines of separase (DEX:ESP-RNAi) expressing PIN2 under its native promoter (Moschou *et al*., 2013)(**Fig. 1**). We observed PIN2 polarity loss in the DEX:ESP-RNAi lines already 12 h after dexamethasone application. Yet, no effect was observed in the corresponding WT plants expressing PIN2 after dexamethasone application (data not shown). These results suggests that the PIN2 localization switch does not depend on temperature and therefore our experimental setup can be used to infer proteolytic targets of separase.

**Figure 1.**
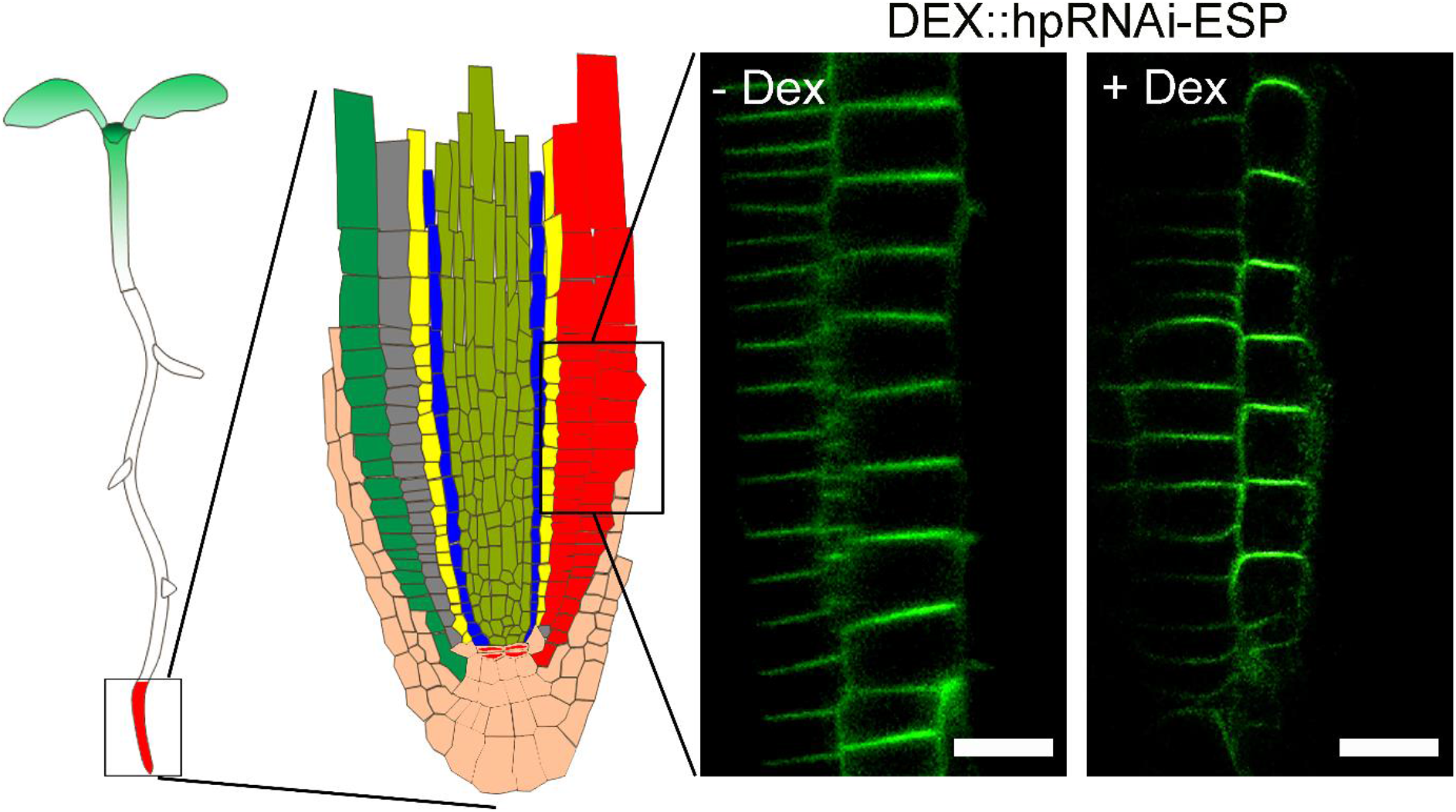
Temperature does not underpin the rootward-to-shootward PIN2 switch in *rsw4*. DEX::hpRNAi-ESP plants expressing PIN2::PIN2-GFP were treated for 12 h with 2 μM dexamethasone (in DMSO) and observed using confocal microscopy. The root region and cells observed (cortex and epidermis) are shown on the left. Representative images of an experiment replicated three times. Scale bars, 5 μm.

### Systematic characterization of root tip N-terminomes

Proteins are always synthesized with methionine (iMet) as the first Nt residue even when a non-canonical translation initiation codon (i.e. non-AUG) is used. However, largely depending on the nature of the second amino acid residue, this first iMet is often cleaved from mature proteins by the N-terminal Met excision (NME) pathway through the Met aminopeptidase (MAP)(Bonissone *et al*., 2013). The N-terminome consists of all peptides that possess mature protein Nt (±iMet) and neo-Nt that are proxies of *in vivo* proteolytic events.

To gain insights into the proteolytic landscape of root cells we analysed the *in vivo* N-terminomes of root tips from 7-d-old seedlings of WT and *rsw4* plants grown at the restrictive temperature for 24 h by a modified version of COFRADIC (**Materials and Methods** and **Fig. 2A, B**). COFRADIC also enables the identification of sequences spanning the scissile peptidic bonds (Vandekerckhove *et al*., 2004; Stes *et al*., 2014). The root tip proteomes of WT and *rsw4* were labelled with N-hydroxysuccinimide esters of either light (WT) or heavy (*rsw4*) isotopic variants of butyric acid to mass tag both Nt arising from translation and neo-Nt. The differential mass tagging of Nt (heavy versus light) enabled us to determine the proteome origin (*rsw4* or WT, respectively) of each labelled peptide.

**Figure 2.**
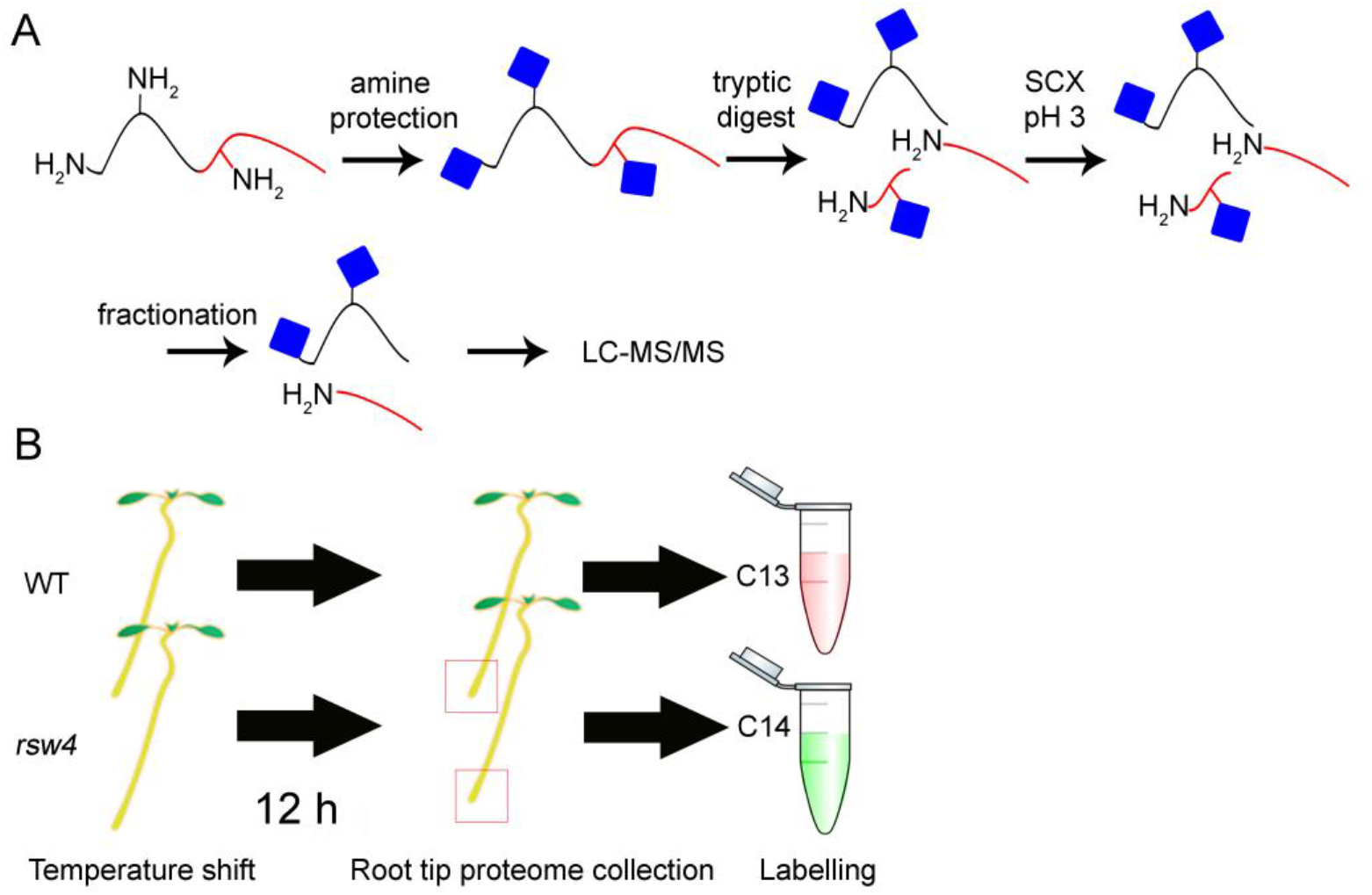
The COFRADIC pipeline and the experimental setup. A. The COFRADIC steps are i) labelling: cysteine alkylation and N-triedeutero-acetylation (butylation); ii) trypsin digestion; ii) pyroQ removal and SCX at low pH. B. WT and *rsw4* seedlings were exposed for 12 h at 28°C and their root tips were harvested, their proteomes extracted and differentially labelled by staple isotopic tags and analysed by COFRADIC (see also Materials and Methods).

To identify translational Nt (peptides containing either the iMet or the 2^nd^ amino acid residue of the protein), we filtered-out peptides carrying neo-Nt (peptides containing Nt beyond the 2^nd^ residue). Out of the total number of 6,974 peptides identified (all present in both genotypes with one exception discussed later), 2,840 possessed translationally yielded Nt converging to 1,057 proteins (**File S1**). Out of 149 unique proteins retaining iMet, 58 proteins had as 2^nd^ residue glutamate (E), 30 aspartate (D) and 20 lysine (K) (E>D>K>>X) (**Fig. 3A**), suggesting that residues with sufficiently large gyration radius block MAP activities (Van Damme *et al*., 2011; Van Damme *et al*., 2012). Protein species devoid of iMet showed strong preference for alanine (A; 29%) and serine (S; 11%) at the 2^nd^ position (P1’; P1^iMet^-P1’-P2’-…-Pn’)(**Fig. 3A**). The functional importance of the 2^nd^ residue can be related to the efficiency of MAP to remove the iMet residue (Shemesh *et al*., 2010). We thus confirm here that A and S residues are among the most preferred ones in the P1’ position of MAP substrates in root tip cells (Frottin *et al*., 2006).

**Figure 3.**
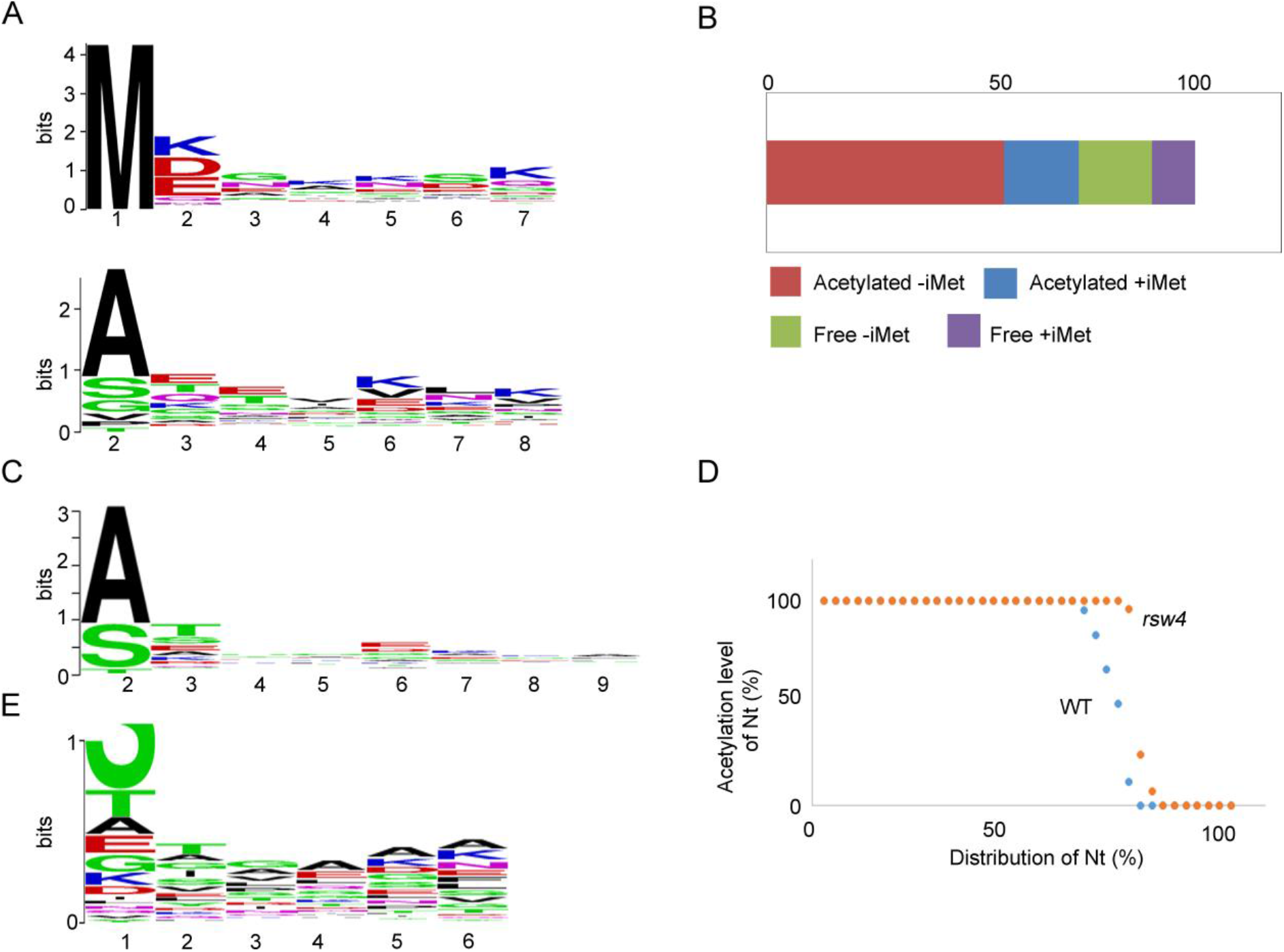
Gene ontology (GO) terms of identified root tip N-termini and residue enrichments at position 2 of proteins with or without N-terminal methionine (iMet). A. Enriched residues of proteins with (top) or without iMet (bottom). B. Distribution (%) of Nt-acetylated peptides lacking iMet (-iMet), with iMet (+iMet) and peptides that were not Nt-acetylated (with or without iMet; -iMet or +iMet, respectively). C. Acetylation specificity of peptides lacking iMet at the Nt (pooled data from WT and *rsw4*). D. Acetylation level of peptides in WT and *rsw4* expressed as a ratio between the acetylated/non-acetylated forms of the same peptides. E. P1’-P6’ specificity of limited proteolysis in root tip cells.

### Acetylation of native Nt in root tips confirms previous studies and adds to the known plastid types capable of N-terminal acetylation

The exposed Nt of proteins is one of the major determinants of protein stability/half-life and certain residues, when exposed at the Nt of a protein, act as triggers for degradation and are therefore called “N-degrons” (Bachmair *et al*., 1986). Nt-acetylation (Nt-ace) can target proteins for degradation via a branch of the N-end rule pathway (Hwang *et al*., 2010). Nt-ace of nascent polypeptides is carried out by N-α-acetyltransferases (NATs) (reviewed in (Gibbs, 2015). Nt-ace typically occurs co-translationally, and, in contrast to lysine (K) acetylation, it seems irreversible. Residues that are acetylated following iMet excision can also act as Nt-ace-degrons when they are not shielded by correct protein folding and/or assembled into oligomeric complexes. In this way, the acetylation-mediated N-end rule plays important roles in general protein quality control.

Between 70 and 90% of eukaryotic proteins carrying Nt-ace and iMet-ace can be targeted for degradation when followed by a bulky residue (e.g. tryptophan; W) (Zhang *et al*., 2015). Out of 2,840 Nt peptides identified 2,067 were *in vivo* acetylated (72%), with 496 carrying iMet-ace (24% of Nt-ace and only 17% of all characterized Nt) (**Fig. 3B**). The majority of experimentally identified Nt-ace (76%) are acetylated after removal of the iMet, displaying at the 2^nd^ position A>S>threonine (T)>>G (**Fig. 3C**). Notably, a similar pattern of acetylation affinity for the 2^nd^ Nt residue has been reported for humans and Arabidopsis leaves (Linster *et al*., 2015). Interestingly, we identified one neo-Nt carrying acetylation. Manual examination of this peptide showed that it corresponds to the mature form of the plastid protein CUTA which carries a peptide leader (Burkhead *et al*., 2003). This observation is consistent with previous results demonstrating acetylation of chloroplastic proteins (Rowland *et al*., 2015), extending the Nt-ace for plastids, as well.

### Nt-acetylation is increased in rsw4

The log2 ratios of light (WT) to heavy (*rsw4*) signals for all identified peptides were calculated with the Mascot Distiller software. The standard deviation of these values for all the peptides (according to the Huber-scale estimator (Foyn *et al*., 2013)) was 0.39 with the median very close to *zero* (0.09; **File S2**). These data suggest that the proteome of *rsw4* shows specific changes rather than pleiotropic redistributions and therefore represents an excellent model for the study of targeted proteomic changes.

In WT, 25 acetylated peptides (from 5 proteins) carrying iMet were significantly enriched (5 proteins; 0.5>log2), in contrast to 61 acetylated peptides (20 proteins) enriched in *rsw4* (**Fig. 3D**). After the removal of the iMet, 1,567 peptides were acetylated of which 105 (19 proteins) were enriched in WT and as many as 359 (20 proteins) enriched in *rsw4*. These data point to an increased accumulation of proteins carrying Nt-ace in *rsw4* root cells, which can be a result of either the enhancement of the acetylation pathway or the abrogation of a proteolytic machinery normally degrading these proteins.

In plants, Nt-ace is indispensable since NatA loss-of-function mutant is embryo lethal (Linster *et al*., 2015). Furthermore, the Nt-ace of SUPPRESSOR OF NPR1, CONSTITUTIVE 1 (SNC1) is important for plant immunity (Xu *et al*., 2015). A specific branch of the N-end rule pathway known as the acetylation N-end rule, targets Nt-ace as degrons and send them for degradation by the proteasome (Liu & Moschou, 2017) and references therein). In yeast, two E3 ligases that recognize Nt-ace-degrons were identified: the ER-associated DOA10/TEB4 and cytosolic NOT4 (Lee *et al*., 2016). Furthermore, a link between anaphase and protein acetylation was suggested in yeast (Van Damme *et al*., 2011).Whether separase crosstalks with the N-end rule pathway and whether Nt-ace impinges on chromosomal segregation in plants merits further investigation. It is not surprising however, that separase-mediated limited proteolysis could crosstalk with digestive proteolytic pathways, as it has been recently discussed (Minina, E.A. *et al*., 2017).

### Identification of neo-Nt peptides in root proteomes and thus potential targets of separase

To identify neo-Nt, i.e. the proxies for proteolytic cleavage, we applied the following filtering criteria: i) positive selection of peptides with butyrylated Nt and ii) exclusion of butyrylated iMet and 2^nd^ residues of the protein. Following filtering, we identified 247 neo-Nt peptides corresponding to 48 unique proteins (**File S1**). The preferable P1’residue of neo-Nt was S, followed by T, A, and E (**Fig. 3E**).

Next, we focused on two classes of peptides: the significantly enriched neo-Nt peptides in WT or *rsw4* (based on the labelling used). As the former might also appear because of differential protein levels in the input proteomes, we scanned the data for the corresponding precursor protein Nt or other neo-Nt peptides from these precursor proteins. We expected to deal with three different scenarios of mass spectroscopy results. First, a specific neo-Nt peptide is present in greater amounts in WT indicating potential separase-mediated (direct or indirect) proteolytic event. Second, a neo-Nt peptide is present in both samples, but in a significantly greater amount in *rsw4*. This scenario would indicate that the corresponding protein was cleaved at the specified site by a protease, which was up/mis-regulated in *rsw4* or that the protein fragment was less stable in WT. Third, the neo-Nt peptide are in equal amounts in both samples, indicating that this cleavage event reflects a general metabolic process not affected by separase depletion.

With the exception of the AT1G79320 (METACASPASE6) for which a neo-Nt was identified only in *rsw4* (removal of a 10 kDa Nt fragment), and may reflect the deregulated expression of this gene, all other neo-Nt-containing peptides were present in both WT and *rsw4* (**File S1**). Nine neo-Nt of proteins were preferentially overrepresented in WT, while four in *rsw4* (**Table 1**). Most of the cleavage events in the identified proteins occurred in coils or α-helixes (**Fig. 4**), a result consistent with previous findings in non-plant organisms (Timmer *et al*., 2009).

**Table 1.** Preferentially cleaved proteins (ratio >2) in WT and *rsw4* root tips. ^1^Only neo-Nt peptides were identified; ^2^Non neo-Nt peptides were significantly increased in *rsw4*.

| Accession | ID | Sequence | Ratio | Function |
| --- | --- | --- | --- | --- |
| ***Preferentially cleaved in WT*** |  |  |  |  |
| AT1G35720 | Annexin 1 | SKAQINATFNR | 75,1 | Calcium-dependent phospholipid binding |
| AT5G63400 | Adenylate kinase 1 | TVTQAEKLDEMLKR | 15,2 | Nucleotide kinase activity |
| AT4G36930 | SPATULA | TTTTTASLIGVHG | 10,1 | Cell division control |
| AT3G05420 | Acyl-CoA binding protein 4 | ATSGPAYPER | 13,0 | Acyl-CoA binding |
| AT2G17720 | Oxygenase<sup>1</sup> | KSETSSGDEEGNGER | 2,1 | Modifies the extensin proteins in root hair cells |
| AT1G80660 | H(+)-ATPase 9<sup>1</sup> | EAQWAQAQR | 2,1 | ATPase, P-type, H+ proton pump |
| AT2G18960 | H(+)-ATPase 1<sup>1</sup> | ELSEIAEQAKR | 2,0 | ATPase, P-type, H+ proton pump |
| AT4G11150 | V-ATP subunit E1 | QDYEKKEKQADVR | 2,0 | ATPase, V1/A1 complex, subunit E |
| AT5G09978 | PROPEP7 | SVVSGNVAAR | 2,0 | Plant immunity |
| ***Preferentially cleaved in rsw4*** |  |  |  |  |
| AT1G13710 | KLU<sup>1</sup> | VLAALAKR | 1031,9 | Positive regulation of cell proliferation |
| AT3G49120 | Peroxidase PRX34<sup>2</sup> | SALVDFDLR | 12,8 | Unidimensional cell expansion |
| AT1G55490 | Chaperonin 60 beta<sup>1</sup> | IVNDGVTVAR | 10,2 | SAR, suppression of protein aggregation. |
| AT3G09740 | Syntaxin of plants 71<sup>1</sup> | ATSDLKNTNVR | 2,3 | protein targeting to membrane, vesicle docking and fusion |

**Figure 4.**
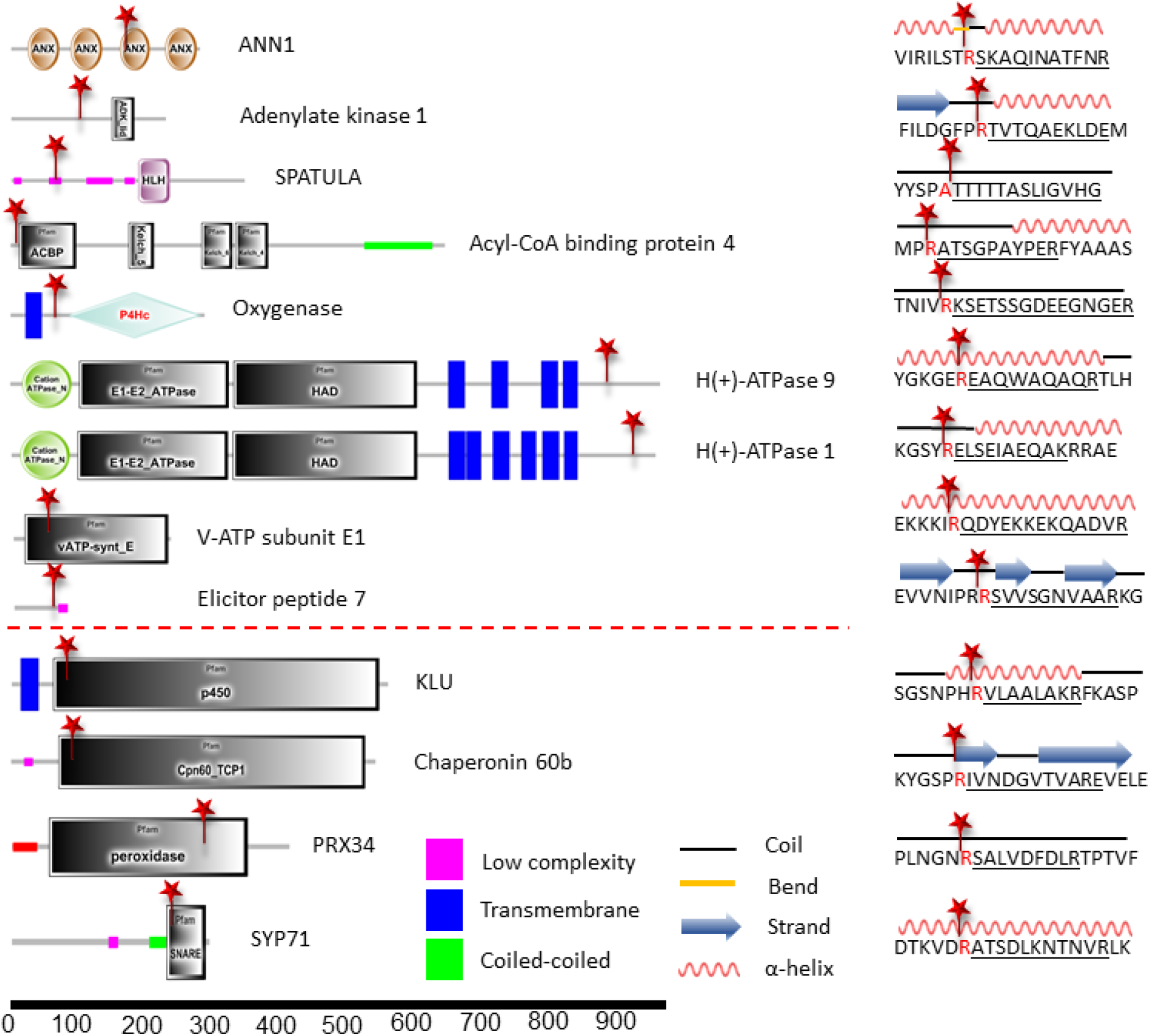
Domain architecture (left), cleavage sites and secondary structures (right) of the neo-N-termini enriched in WT (above the dotted red line) or in *rsw4* (below the dotted line). Domain arcitecture was defined by SMART (Letunic *et al*., 2015). Secondary structures represent predictions defined by YASPIN (Lin *et al*., 2005). The only proteins identified from these cleaved in loops are the V-ATPase E1 and PRX34.

Interestingly, neo-Nt corresponding to the V-ATPase E1, H+-ATPases, Annexin 1 (increased in WT) and the SYNTAXIN 71 (SYP71; increased in *rsw4*) were the most striking proteins relevant to the vesicular trafficking and auxin defects observed in *rsw4*. Annexin 1 and H+-ATPases, are plasma membrane proteins (Baucher *et al*., 2011; Haruta *et al*., 2015), while the V-ATPase E1 localizes at the tonoplast and the TGN (Strompen *et al*., 2005; Schumacher & Krebs, 2010). Interestingly, separase localizes at the plasma membrane and TGN at least transiently (Moschou *et al*., 2013). We should note that for the two H+-ATPases (as well as the identified oxygenase) we could not ascribe additional peptides except the identified neo-Nt and therefore, they could be simply more abundant in WT.

In plants, V-ATPase activity acidifies the TGN and the vacuole (Strompen *et al*., 2005; Schumacher & Krebs, 2010). In mammalian cells, separase depletion leads to a decrease in constitutive protein secretion as well as endosomal receptor recycling and degradation (Bacac *et al*., 2011). Depletion of separase results in increased vesicular pH mediated by V-ATPase inhibition. It is thus reasonable to assume that plant separase may also regulate the acidification of vesicles by modulating the V-ATPase activity.

The H+-ATPases (AHA) regulate cellular acidification and cell expansion by building a proton gradient across the plasma membrane. Downregulation of AHA activity occurs after treatment of plants with the growth inhibitory peptide, RALF, leading to the inhibition of cell expansion (Haruta *et al*., 2014). Processing might be a complementary mechanism to control the activity of AHA proteins. Plants with a hyperactive allele of AHA2 (*ost2*) show abscisic acid insensitivity and drought sensitivity (Merlot *et al*., 2007). Furthermore, evidence suggest a crosstalk between auxin signalling and acidification that involves the activation of AHA in Arabidopsis roots (Fendrych *et al*., 2016). A more complete understanding of hprocessing may control AHA activity will provide new prospects for improving plant growth and drought tolerance.

Annexin 1 (ANN1) is a root-specific annexin regulating root length via sensing pH-dependent channel activity (Gorecka et al., 2007). In pea (*Pisum sativum*) an ANN1 homologue can also redistribute in response to gravistimulation (Clark et al., 2000). In maize cells, other annexins are known to stimulate Ca^2+^-dependent exocytosis (Carroll *et al*., 1998). In animal cells, cleavage of ANN1 at the N-terminus by calpain protease mediates the secretory processes by modifying either requirements for Ca^2+^ or ANN1 efficacy in membrane binding, vesicle aggregation or endocytic processing (Barnes & Gomes, 2002; Sugimoto *et al*., 2016). Other annexins, e.g. A2, is a Ca^2+^-, actin-, and lipid-binding protein mediating the formation of lipid microdomains required for the structural and spatial organization of fusion sites at the plasma membrane (Gabel *et al*., 2015). It is tempting to speculate that separase may regulate the activity of ANN1 by modifying its requirement for Ca^2+^. Interestingly, the Arabidopsis separase has an EF-hand, responsible for Ca^2+^ binding.

The identified neo-Nt of the cytoplasmic Acyl-CoA-binding protein 4 (ACBP4) is preferentially enriched in WT (**Table 1**). ACBP4 is involved in the transport of lipids (C ≥20) to the plasma membrane for the biosynthesis of surface lipids such as wax and cutin (Du *et al*., 2016). Depletion of AtACBP4 in Arabidopsis resulted in decreases in galactolipids and phospholipids, suggesting its role in membrane lipid biosynthesis, as well as diminished capability in the generation of signals such as salicylate for induction of systemic acquired resistance. It is yet unclear what could be the role of ACBP4 processing but may lead to protein activity modulation. A reduced activity of ACBP4 in *rsw4* could compromise lipid signalling that could contribute to the observed phenotypes.

The product of PROPEP7 (**Table 1**; enriched in WT) is a member of the PROPEP family of Damage Associated Molecular Pattern (DAMP) elicitor peptides. The cleavage on the PROPEP7 is interesting, since it corresponds to the predicted cleavage site that will release the Pep7 elicitor peptide, but has never been experimentally shown (Bartels & Boller, 2015). Furthermore, this family is believed to have extended functionalities beyond wounding and immunity, potentially in growth and development (Bartels & Boller, 2015). In fact, overexpression of PROPEP1 leads to larger root system in Arabidopsis (Huffaker *et al*., 2006). Interestingly, the identified cleavage event enriched in *rsw4* on Chaperonin 60 beta may also be involved in pathogen resistance (**Table 1**). The corresponding mutant, *Arabidopsis lesion initiation 1* (*len1*), develops necrotic lesions which may activate systemic acquired resistance (Ishikawa *et al*., 2003), without pathogen attack (Ishikawa, 2005).

The peroxidase PRX34 was found at significantly higher levels in *rsw4* (both neo-Nt and internal peptides). PRX34 is an apoplastic reactive oxygen species (ROS) generator involved in resistance to pathogens in Arabidopsis (Daudi *et al*., 2012) and also root development through a genetic interaction with the mitogen-activated protein kinase 6 (MAPK6) (Han *et al*., 2015). Interestingly, *PRX34* mutants show larger leaves than WT. Similarly, oxidases in the apoplast of tobacco plants modulate resistance to pathogens and plant development in a pathway that involves ROS-MAPKs (Moschou *et al*., 2008; Moschou *et al*., 2009; Tisi *et al*., 2011; Gemes *et al*., 2016). We assume that it is unlikely that PRX34 is a direct separase target considering PRX34 localization in the apoplast, but its deregulation is rather an indirect effect.

The Qb-SNARE SYP71 (enriched in *rsw4*) is part of a bigger family of proteins mediating vesicular fusion (El Kasmi *et al*., 2013). Together with Qb-SNARE NPSN11 and VAMP721,722 the SYP71 forms a tetrameric KNOLLE-containing complex. KNOLLE mediates vesicular trafficking during Arabidopsis cytokinesis and its targeting to the cytokinetic apparatus known as cell plate is compromised in *rsw4* (Moschou *et al*., 2013). Hence, a plausible scenario is that SYP71 is preferentially degraded in *rsw4*, an event which may impact vesicular fusion. It is highly likely that vesicular fusion is compromised in the absence of separase as it has been shown in Arabidopsis (Moschou *et al*., 2013) and *C. elegans* (Bembenek *et al*., 2010; Mitchell *et al*., 2014).

Finally, SPATULA is a negative (neo-Nt increased in WT), while KLU (neo-Nt increased in *rsw4*) a positive cell division regulator (Alvarez & Smyth, 1999; Heisler *et al*., 2001; Anastasiou *et al*., 2007; Ichihashi *et al*., 2010; Josse *et al*., 2011; Makkena & Lamb, 2013). SPATULA is a transcription factor required for specification of carpel and valve tissues by restricting cell division. The corresponding mutants show enhanced root and leaf growth. KLU produces an as yet unkown mobile signal with cell division promoting properties. KLU is involved in generating a mobile growth signal distinct from the classical phytohormones that defines primordium size.

### Consensus cleavage sites of substrates enriched in wild-type amd thus potential separase targets

Using iceLogo, we inspected the amino acid residues at the prime (P’) and nonprime (P) substrate positions and analysed the frequencies of specific residues after statistical correction by means of the natural occurrence of amino acids in Arabidopsis proteins (Colaert *et al*., 2009). Unlike what has been described for kleisin motifs cleaved by separase, our data do not show specificity of Arabidopsis separase against EXXR (X, any residue) sequence (Sullivan *et al*., 2004; Lin *et al*., 2016) (**Fig. 5**). This finding suggests that either not all identified substrates are direct targets of separase or that separase in plants has evolved broader cleavage specificity. We should note that so far sequence specificity for separase in non-plants has been deduced from cleavage sites on only few proteins related to cell division, i.e. SLK19 and kleisins and may not reflect the general specificity of separases. Noteworthy, the kleisins in plants are a multimember gene family (da Costa-Nunes *et al*., 2006; Ma *et al*., 2016; Minina, E. A. *et al*., 2017). Hence, plant separases may have adapted to the expansion of plant kleisins by adjusting accordingly their substrate recognition repertoire.

**Figure 5.**
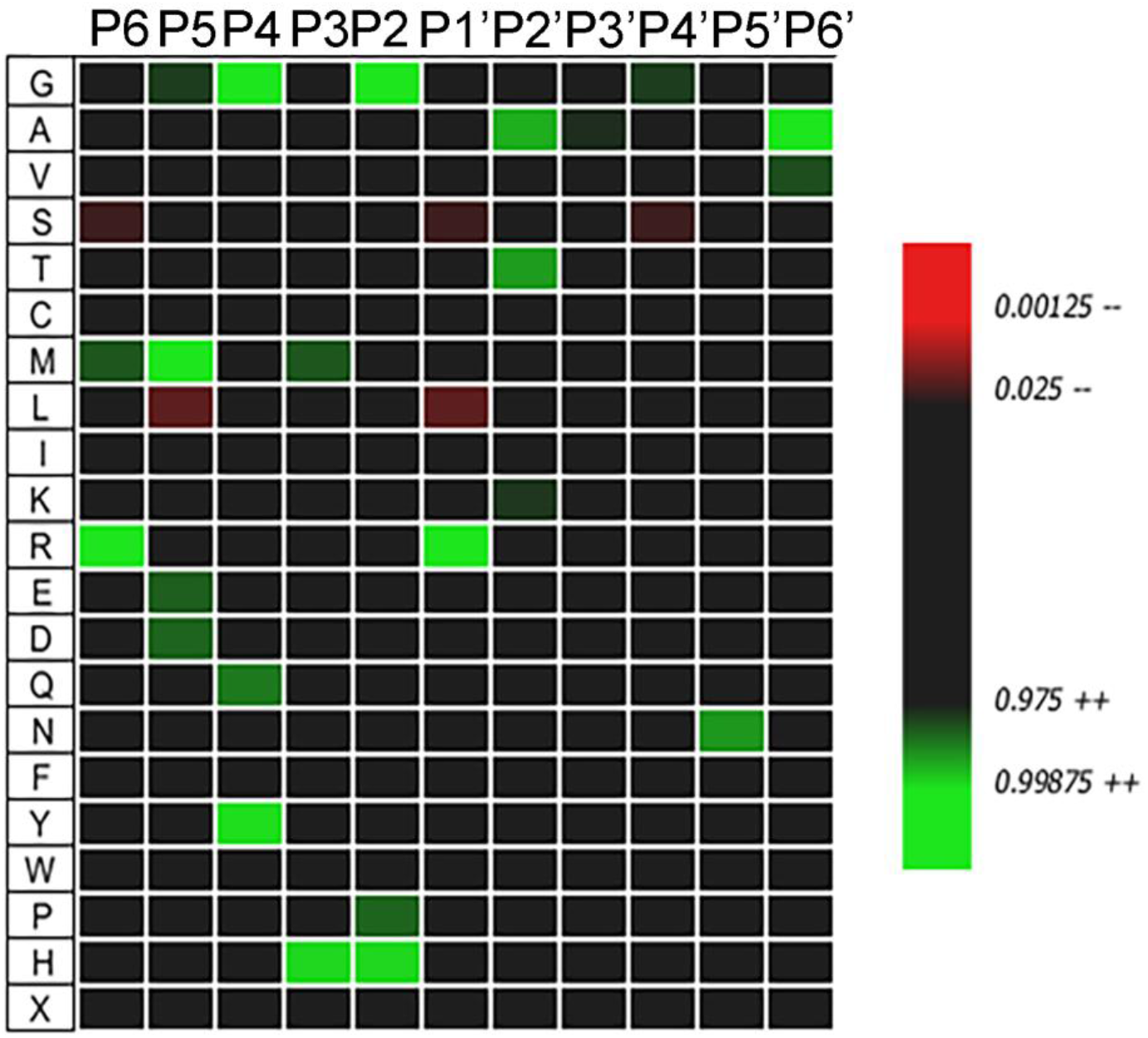
Icelogo showing the specificity of cleavage events enriched in WT.

## Conclusion

We previously established a role for separase in controlling vesicular trafficking and plant development (Moschou *et al*., 2016a). Likewise, *C. elegans* separase is required for secretion of Rab11-positive vesicles, and in human cells separase controls fusion and acidification of vesicles (Bembenek *et al*., 2010; Bacac *et al*., 2011; Mitchell *et al*., 2014). Interestingly, fusion of vacuolar membranes in yeast requires proteasomal degradation of ubiquitinated Ypt7, a yeast homolog of Rab7 GTPase (Kleijnen et al. 2007). Ubiquitination may also regulate membrane fusion events that reassemble fragmented organelles after mitosis in mammalian cells and vacuole-plasma membrane fusions in plant cells during plant immunity (Meyer & Popp, 2008). Yet, the role of processing in vesicular trafficking remains unknown. Here, we show that plant separase may be involved in the regulation of acidification and lipid signalling, and we assign potential molecular targets for this function. Furthermore, we show that the proteolytic landscape of *rsw4* mutants involves proteins that modulate resistance to pathogens, cell expansion and cell division. This finding extents the possible functions of separase, suggesting potential links of a core component of cell division and development to plant immune responses.

## Materials and Methods

### Plant Material and Growth Conditions

*Arabidopsis thaliana* wild-type (WT) and *rsw4* plants in the Col-0 ecotype background were grown on vertical plates containing half-strength Murashige and Skoog (MS) medium supplemented with 1% (w/v) sucrose and 0.7% (w/v) plant agar, at 20°C (permissive temperature) or 28°C (restrictive temperature), 16/8-h light/dark cycle, and light intensity 150 μE m^−2^ s^−1^. The root swelling phenotype was observed in homozygous *rsw4* plants within 3 d of incubation at 28°C. The dexamethasone RNAi lines are in Col-0 background and have been previously described (Moschou *et al*., 2013).

### A shortened N-terminal COFRADIC protocol: sample preparation

Seedlings grown on MS plates with nylon filters (0.8 μm) were harvested and root tips were collected after manual excision with a blade. Tips were snap frozen in liquid N2, and ground into a fine powder with a mortar and pestle. Samples were prepared as previously described (Tsiatsiani *et al*., 2014). To achieve a total protein content of 1 mg, 0.2 g frozen ground tissue was re-suspended in 1 mL of buffer containing 1% (w/v) 3-[(3-cholamidopropyl)-dimethylammonio]-1-propanesulfonate (CHAPS), 0.5% (w/v) deoxycholate, 5 mM ethylenediaminetetraacetic acid, and 10% glycerol in 50 mM HEPES buffer, pH 7.5, further containing the suggested amount of protease inhibitors (one tablet/10 mL buffer) according to the manufacturer’s instructions (Roche Applied Science). The sample was centrifuged at 16,000g for 10 min at 4°C, and guanidinium hydrochloride was added to the cleared supernatant to reach a final concentration of 4 M. Protein concentrations were measured with the DC protein assay (Bio-Rad), and protein extracts were further modified for N-terminal COFRADIC analysis as described previously (Staes et al., 2011).

Col-0 (WT) primary amines were labelled with the N-hydroxysuccinimide (NHS) ester of ^12^C_4_-butyrate and *rsw4* with NHS-^13^C_4_-butyrate, resulting in a mass difference of approximately 4 Da between light (^12^C_4_) and heavy (^13^C_4_) labelled peptides. After equal amounts of the labelled proteomes had been mixed, tryptic digestion generated internal, non-N-terminal peptides that were removed by strong cation exchange at low pH (Staes *et al*., 2011). Due to the low amount of input material, the COFRADIC protocol was cut short after the first reversed phase-high performance liquid chromatography (RP-HPLC) step and the resulting 15 fractions were subjected immediately for identification by LC-MS/MS.

### Peptide Identification, and Quantification

Samples were dissolved in 2% ACN, 0.1% TFA and subjected to ESI-MS/MS on a Q Exactive Orbitrap mass spectrometer operated similarly as previously described with some modifications (Stes *et al*., 2014). Peptides were separated on a reverse phase column with a linear gradient from 98% solvent A’ (0.1% formic acid in water) to 40% solvent B’ (0.1% formic acid in water/acetonitrile, 20:80 (v/v) in 127 min at a flow rate of 300 nL/min. MS was run in data-dependent, positive ionization mode, with the initial MS1 scan (400–2000 m/z; AGC target of 3 × 10^^^6 ions; maximum ion injection time of 80 ms) acquired at a resolution of 70 000 (at 200 m/z), which was followed by up to 10 tandem MS scans of the most abundant ions at a resolution of 17 500 (at 200 m/z) according the following criteria: AGC target of 5 × 10^^^4 ions; maximum ion injection time of 120 ms; isolation window of 2.0 m/z; fixed first mass of 140 m/z; underfill ratio of 1.2%; intensity threshold of 5 × 10^^^3; exclusion of unassigned, 1, 5–8, >8 charged precursors; peptide match preferred; exclude isotopes, on; dynamic exclusion time, 20 s. From the MS/MS data, Mascot Generic Files (mgf) were created using the Mascot Distiller software (version 2.5.1.0, Matrix Science). Peak lists were then searched using the Mascot search engine with the Mascot Daemon interface (version 2.5.1, Matrix Science). Spectra were searched against the TAIR10 database. Mass tolerance on precursor ions was set to 10 ppm (with Mascot’s C13 option set to 1), and on fragment ions to 20 mmu. The instrument setting was on ESI-QUAD. Endoproteinase semi-Arg C/P was used with one missed cleavage allowance, rather than trypsin, beause cleavage after lysine residues is abolished by side chain acylation. Variable modifications were set to pyroglutamate formation of N-terminal glutamine and fixed modifications included methionine oxidation to its sulfoxide derivative, S-carbamidomethylation of cysteine, and butyrylation (^12^C_4_ or ^13^C_4_) of the lysine side chain and peptide N-termini. Peptides with a score higher than the MASCOT identity threshold set at 99% confidence were withheld. The Mascot Distiller Toolbox (version 2.5.1.0; Matrix Science) was used to quantify relative peptide abundance and in case of doubt were manually curated (for example for singletons). MS data have been deposited to the ProteomeXchange Consortium via the PRIDE (Vizcaino *et al*., 2016).

## Supplemental Data

**Supplemental File 1**. N-terminome of WT and *rsw4* Arabidopsis root tips.

**Supplemental File 2**. Variance distribution of peptides quantifications in WT and *rsw4*.

